# Acute increase of protein O-GlcNAcylation in mice leads to transcriptome changes in the brain opposite to what is observed in Alzheimer’s Disease

**DOI:** 10.1101/2024.09.19.613769

**Authors:** Margaret Bell, Mariame S Kane, Xiaosen Ouyang, Martin E Young, Anil G Jegga, John C. Chatham, Victor Darley-Usmar, Jianhua Zhang

## Abstract

Enhancing protein O-GlcNAcylation by pharmacological inhibition of the enzyme O-GlcNAcase (OGA) is explored as a strategy to decrease tau and amyloid-beta phosphorylation, aggregation, and pathology in Alzheimer’s disease (AD). There is still more to be learned about the impact of enhancing global protein O-GlcNAcylation, which is important for understanding the mechanistic path of using OGA inhibition to treat AD. In this study, we investigated the acute effect of pharmacologically increasing O-GlcNAc levels, using OGA inhibitor Thiamet G (TG), on normal mouse brains. We hypothesized that the transcritome signature in respones to TG treatment provides a comprehensive view of the effect of OGA inhibition. We sacrificed the mice and dissected their brains after 3 hours of saline or 50 mg/kg TG treatment, and then performed mRNA sequencing using NovaSeq PE 150 (n=5 each group). We identified 1,234 significant differentially expressed genes with TG versus saline treatment. Functional enrichment analysis of the upregulated genes identified several upregulated pathways, including genes normally down in AD. Among the downregulated pathways were the cell adhesion pathway as well as genes normally up in AD and aging. When comparing acute to chronic TG treatment, protein autophosphorylation and kinase activity pathways were upregulated, whereas cell adhesion and astrocyte markers were downregulated in both datasets. Interestingly, mitochondrial genes and genes normally down in AD were up in acute treatment and down in chronic treatment. Data from this analysis will enable the evaluation of the mechanisms underlying the potential benefits of OGA inhibition in the treatment of AD. In particular, although OGA inhibitors are promising to treat AD, their downstream chronic effects related to bioenergetics may be a limiting factor.

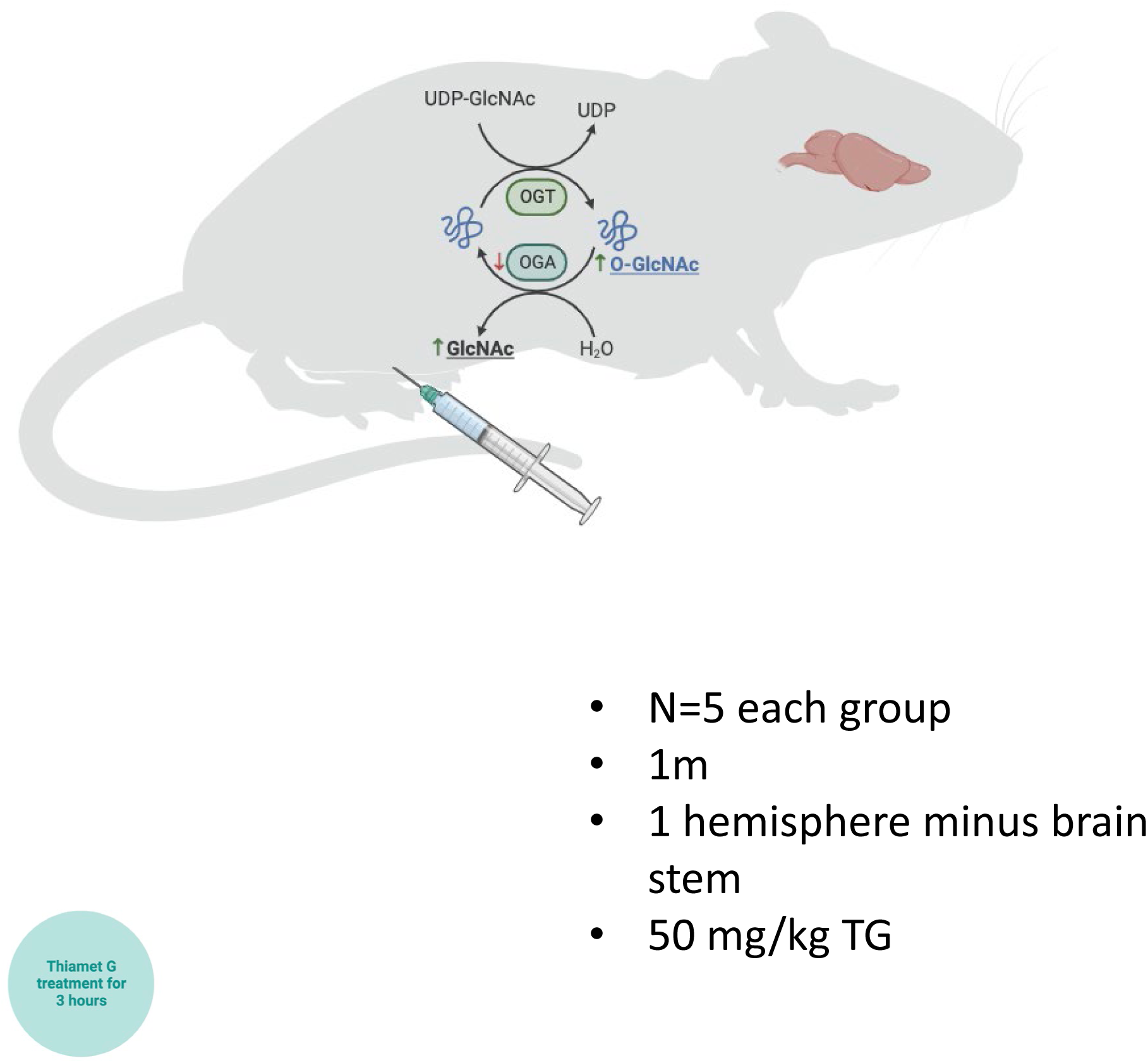

## INTRODUCTION

Protein O-GlcNAcylation is a post-translational modification by O-linked β-N-acetylglucosamine (O-GlcNAc)^1,2^. O-GlcNAc is added to hydroxyl groups of serine/threonine residues of proteins by the enzyme O-GlcNAc transferase (OGT) and removed by the enzyme O-GlcNAcase (OGA)^3,4^. O-GlcNAc has been identified on proteins localized in essentially all cellular compartments, including the nucleus, membrane, cytoplasm, and mitochondria^2,5^. O-GlcNAc regulates biological processes such as cell signaling and energy metabolism, by changing the activities of downstream effector molecules^2,5–7^. In addition, O-GlcNAc plays a role in gene and protein expression by binding to proteins responsible for modulating critical steps in transcription and translation^2,7–9^. In humans, mutations in OGT are associated with intellectual disabilities^10,11^, while embryonic deletion of OGA or OGT results in lethality in mice^12,13^; thus the dynamic regulation of O-GlcNAcylation is important in maintaining cell survival, function and homeostasis.

O-GlcNAc modification has been observed on many key proteins involved in the pathogenesis of various neurodegenerative diseases^14^. Alzheimer’s disease (AD) is the most prevalent neurodegenerative disease, causing cognitive impairment and memory loss^15^. One pathological hallmark of AD is the hyperphosphorylation of the tau protein and the formation of neurofibrillary tangles^16–18^. Controversial results have been reported regarding how O-GlcNAc abundance regulates tau protein hyperphosphorylation in AD. Liu et al showed that O-GlcNAc abundance negatively regulated tau phosphorylation *in vivo* and *in vitro* due to both modifications binding to similar sites (serine/threonine)^19^. On the other hand, a mass spectrometry study comparing a transgenic model of AD to wildtype mice revealed that there was only one O-GlcNAc site on the tau protein (S400) in wildtype mice which was undetected in AD, contesting the hypothesis that O-GlcNAc abundance can block tau hyperphosphorylation^20^. Similar findings have shown both an increase^21^ and decrease^22^ of total protein O-GlcNAcylation in AD postmortem samples. However this controversy has been reconciled to some extent. A proteomic study by Wang et al showed decreased O-GlcNAc in a small sub-proteome and increased in a larger sub-proteome in AD patients^23^. The exact role of O-GlcNAc in AD is not known, however the consistent presence of O-GlcNAc on key proteins involved in the pathogenesis of these diseases makes this modification a potential target for treatment development.

Studies investigating the effects of changes in O-GlcNAc levels on neurodegenerative pathologies have also shown conflicting results. On one hand, the use of OGA inhibitors in mouse models of AD show benefits such as decreased tau phosphorylation in tau overexpression models^24,25^. This has led to multiple OGA inhibitors being tested in AD clinical trials, however the exact mechanism by which they affect AD pathology is unclear^26^. On the other hand, work performed using OGA loss of function in a *C. elegans* neurodegenerative model shows that it increases proteotoxicity^27^. Our lab treated mice with a potent and highly specific OGA inhibitor, Thiamet G (TG) for 3 hours to analyze the proteomic profile of the brain after increased O-GlcNAc expression^28^. We identified 85 peptides that were significantly altered by acute TG treatment (fold change greater than 1.5)^28^. Additional studies by Taylor et al showed that acute inhibition of OGA leads to reduced learning and memory, as measured using a novel recognition test^29^. Furthermore, we assessed chronic TG treatment in mice for 2.5 months^30^. We performed behavioral tests during the TG treatment and RNA sequencing at the conclusion of the treatment. We found that TG treatment improved working memory in mice by Y-maze test and that the top differentially expressed genes (DEG) were related to learning and cognition. Additionally, we uncovered that many transcripts involved in oxidative phosphorylation were downregulated. Although O-GlcNAcylation is known to regulate transcription^31–33^, the effect of acute elevation of O-GlcNAcylation on transcriptomes in the brain is unknown. In this study, we investigated the acute increase of O-GlcNAcylation on the mouse brain transcriptome. In addition, we compared transcriptome of acute increase of O-GlcNAcylation with chronic increase of O-GlcNAcylation, which we previoulsy published^30^.

## MATERIALS AND METHODS

### Mice

All mice work was performed with approval by the University of Alabama at Birmingham (UAB) Institutional Animal Care and Use Committee. C57BL/6J (RRID:IMSR_ JAX:000664) mice were bred in house. Mice were housed with access to food and water ad libitum. At 1 month of age, mice were treated intraperitoneally with saline or 50mg/kg of TG to increase overall O-GlcNAc expression (n=5 male mice each group). TG binds OGA with a Ki at low nM range, crosses the blood-brain barrier and is with 37,000-fold selectivity over other glycoside hydrolases^34,35^. Previous studies from our lab show acute treatment of 10 mg/kg TG is sufficient to increase protein O-GlcNAcylation in the mouse brain. 3 hours post-injection (TG injected at 7 am, and mice sacrificed at 10 am), mice were sacrificed using Fatal-Plus (pentobarbital, obtained through UAB Animal Resource Program). Mice were perfused with 1xPBS for 5 mins to remove excess blood. The whole brain was dissected. The brain stem and occipital bulb were removed, and one brain hemisphere was flash frozen in liquid nitrogen for processing. The samples were powdered to obtain a homogeneous brain sample. The brain powder was aliquoted for western blot and RNA sequencing experiments (**Figure 1**).

**Figure 1:**
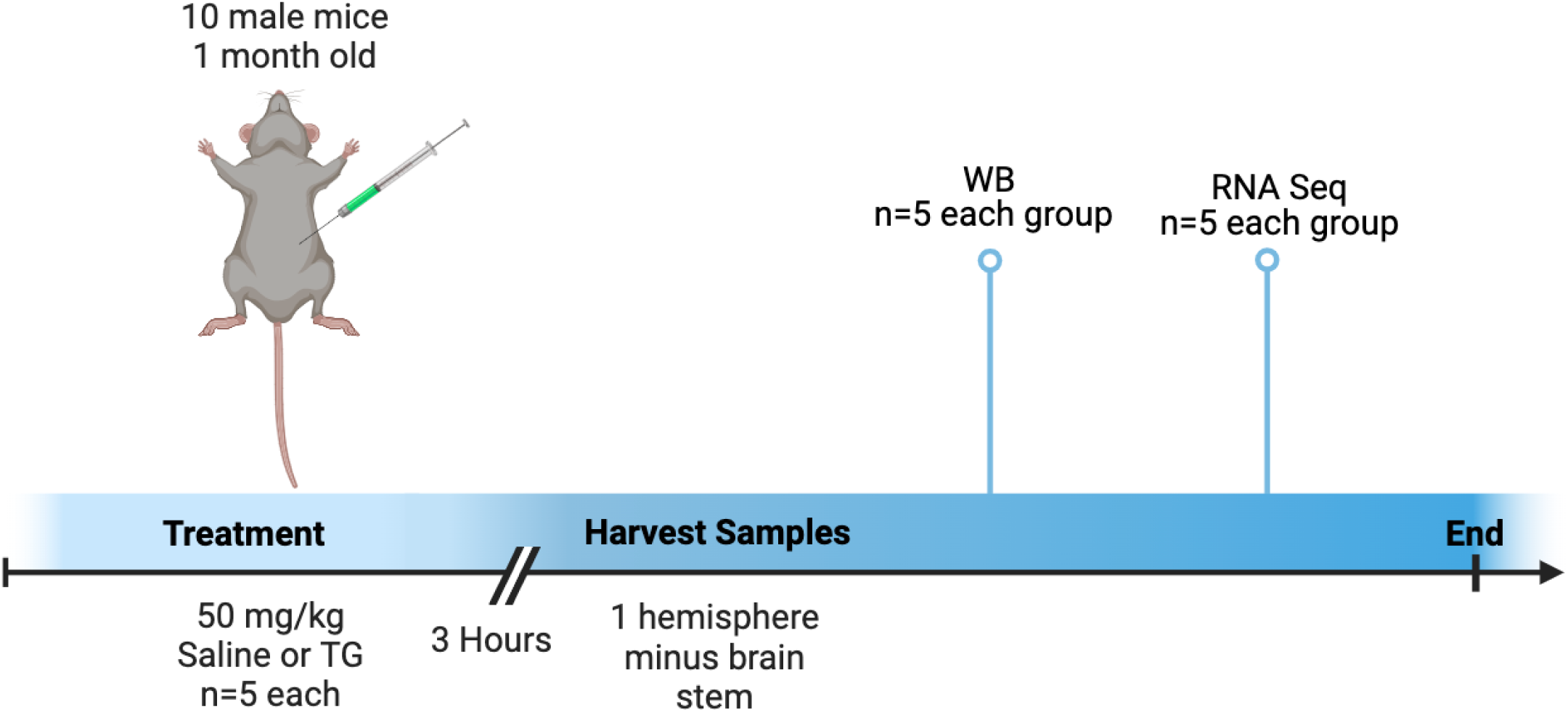
Experimental design flow chart. Ten 1-month old male mice were injected with 50 mg/kg saline or TG. After 3 hours, animals were sacrificed, and one hemisphere of the brain was harvested for downstream analysis. Downstream analysis included western blot and rna sequencing.

### Western Blot

Western blot analysis was performed with flash frozen brain tissue. Protein isolation was performed using modified RIPA buffer with phosphatase (Fisher 1862495) and protease (Sigma #11836170001) inhibitors. Samples were homogenized using bead mill homogenizer on speed 4 for 45 seconds and incubated at room temperature from 10 minutes. Samples were then centrifuged for 10 mins at 4°C and 1000 x g. The supernatant was removed as the homogenized protein sample.

To measure overall levels of O-GlcNAcylated proteins, 25 μg of protein was transferred to 0.45 μm PVDF membrane (ThermoFisher 88518) and probed overnight at 4°C using the primary antibody mouse IgM anti-CTD110.6 (Millipore MABS1254) at 1:2000 in filtered PBS with 1% casein (Biorad 1610783). Goat-anti-mouse IgM secondary antibody (CalBiochem cat# 401225) at 1:10000 in PBS with 1% casein was then used for 2 hours at 4°C. For protein loading control, amido black (Sigma cat# N3393) was used.

To measure OGT protein level, 10 μg of protein was transferred to 0.45 μm PVDF membrane and probed with rabbit anti-OGT IgG (Cell Signaling, #24083; RRID:AB_2716710) primary antibody at 1:1500 in 5% milk overnight at 4°C. Donkey anti-rabbit IgG-HRP (Santa Cruz cat# NA934V) at 1:5000 was used as a secondary antibody. Ponceau (Tocris #5225) was used for protein loading detection.

For OGA, 5 μg of protein was transferred to 0.45 μm PVDF membrane and detected using the Mgea5 primary antibody (Proteintech Cat# 14711-1-AP; RRID:AB_2143063)) at 1:1000 in 5% milk overnight at 4°C. Again, donkey anti-rabbit IgG-HRP (Santa Cruz cat# NA934V) at 1:5000 was used as a secondary antibody for 1 hour at room temperature. Ponceau was used for protein loading control.

All images were quantified using ImageQuant TL v8.1.0.0 (RRID:SCR_018374). For membranes with a clear band, 1D gel analysis with rolling ball or minimum background method was used. For full lane intensity analysis, ImageQuant TL v8.1.0.0 analysis toolbox was used.

### RNA preparation

RNA was isolated from saline or TG treated mice to perform RNA sequencing. Isolation was performed using Qiagen RNeasy Plus Mini Kit (Qiagen cat# 74134). RNA concentration was measured using Qubit 4 Fluorometer (RRID:SCR_018095. For sample quality control, only samples with a concentration ≥20ng/μL, RNA integrity number ≥4, OD260/280 ≥2, and OD260/230 ≥2.

### RNA sequencing

Poly-A mRNA sequencing was performed using Novogene Corporation Inc. Sequencing platform used was Illumina NovaSeq 6000 (RRID:SCR_016387) and sequencing strategy was paired end 150bp acquiring 6G of data per sample. Data quality control was performed by filtering reads containing adapters or with low quality measured by uncertain nucleotides >10% or Qscore of over 50% bases below 5. Reads were then mapped to the mouse reference genome (GRCm38.p6) using hisat2 v2.0.5 (RRID:SCR_015530) and assembled using StringTie v1.3.3b (RRID:SCR_016323). Gene expression was quantified used featureCounts v1.5.0-p3 (RRID:SCR_012919) and a p-value <0.05 threshold was used to further filter the transcripts. Differential gene expression was performed using DESeq2 v1.20.0 (RRID:SCR_015687) to identify differentially expressed between saline and TG groups. Functional enrichment pathway analysis was performed using the ToppFunn or ToppCluster application of the ToppGene Suite (RRID:SCR_005726). Pathways of interest were identified using adjusted p-value <0.05. Networks were visualized using Cytoscape (RRID:SCR_003032) and heatmaps were generated using the Morpheus application (RRID:SCR_017386).

### Statistical analysis

G*Power 3.1 (RRID:SCR_013726) was used for power analysis. The effect size was calculated from previous literature showing an increase in O-GlcNac after 3 h of TG treatment in the mouse brain^36^. Based on this data, the effect size d = 1.84 and the total number of samples needed in order for a statistical power >0.7 is 10. Western blot images were analyzed using ImageQuant TL (RRID:SCR_018374). Graphpad Prism 8.2.1 (RRID:SCR_002798) was used for statistical analysis (normality, outlier, and t-test). Normality was measured using Shapiro-Wilk normality test. Outliers were identified using the ROUT method with Q = 1%. Unpaired t-test was used to analyze normally distributed data, whereas Mann Whitney test was used for non-normal data. Protein and fpkm expressions were significant if p-value < 0.05 and trending towards significance if p-value < 0.1. For RNA sequencing analysis, no log2 FoldChange cutoff was applied. Differential expressed genes were significant if p-value is < 0.05. Pathway enrichment analysis used a cutoff of adjusted p-value < 0.05 (FDR Benjamini & Hochberg correction)

## RESULTS

### Overall protein O-GlcNAcylation is elevated after 3-hour TG treatment

To examine the impact of acute inhibition of OGA, 1 month old male mice were injected with 50 mg/kg of TG (intraperitoneal) or saline for 3 hours before preparation of samples for analysis (**Figure 1**). As shown in **Figure 2A**, O-GlcNAcylated protein levels were increased after TG treatment (unpaired t-test. t=5.948, df=8, p=0.0003), as expected and consistent with the inhibition of OGA^30,36^. The protein levels of the enzymes controlling the amount of protein O-GlcNAcylation, OGA, and OGT, were not altered by TG treatment at this early time point (**Figure 2B,2C**) (OGA: Mann Whitney test. U=9, df=8, p=0.5476) (OGT: unpaired t-test. t=1.259, df=8, p=0.2435).

**Figure 2:**
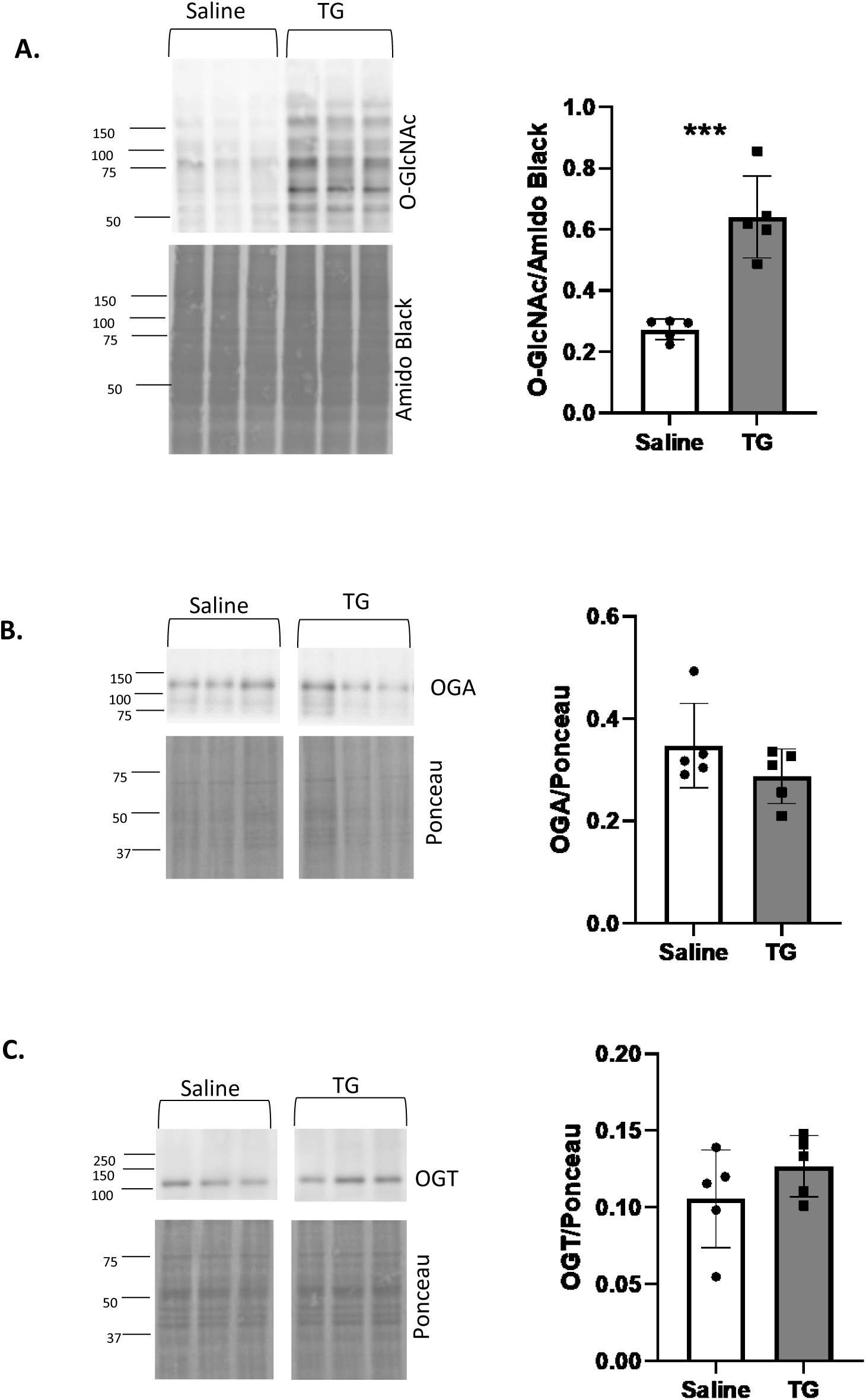
TG treatment increased mouse brain O-GlcNAc protein levels. Mice were sacrificed after 3 hours of TG treatment and O-GlcNAcylated protein levels and the amounts of OGA and OGT determined. **A)** Western blot analysis of overall protein O-GlcNAcylation in the TG treated and control group as assessed using the CTD 110.6 antibody. Amido black staining was used as a loading control. **B)** OGA western blot analysis using the Mgea5 primary antibody from Proteintech with normalization to total protein using Ponceau protein staining. **C)** OGT western blot analysis using the Cell Signaling antibody. All images were analyzed using ImageQuant. All data was analyzed by unpaired Student’s t-test or Mann Whitney test, ***p < 0.001. n=5 mice per group.

### 3-hour TG treatment significantly changes transcriptomes in the brain

Next, we performed RNA sequencing of mouse brain tissue with saline or TG treatment. The overall RNA sequencing analysis workflow is in **Figure 3**. Over 6G of clean base reads were measured for each sample (**Supplemental Table 1**) with >96% mapping to the mouse reference genome (**Supplemental Table 2**). Additional quality control measurements can be found in **Supplemental table 1 and 2**. After filtering for genes with low expression, RNA sequencing detected 27,714 transcripts (**Figure 3, Supplemental Table 3**). When applying a criterion of p-value <0.05, we detected 1,234 transcripts which were significantly differentially expressed in response to TG treatment (**Figure 3, Supplemental Table 4**). As shown in the volcano plot of the 27,714 transcripts, 593 were upregulated and 641 were downregulated (**Figure 4A**). The heatmap of this data presented in **Figure 4B** again shows clear differences in mRNA in response to TG treatment, and the hierarchical clustering segregates by treatment group showing the high integrity of the data. To determine whether there are adaptive / compensatory changes of OGA and OGT mRNA in response to OGA inhibition by TG, we plotted OGA mRNA levels. Indeed, although the protein levels of OGA and OGT did not differ after TG treatment, the OGA transcript exhibited a significant increase in response to TG treatment (Mgea5; Log2Fold Change=0.3101 p=0.0003) (**Figure 4C**), whereas the OGT-normalized mRNA count showed no change (Log2Fold Change=-0.1744 p=0.1980) (**Figure 4D**).

**Figure 3:**
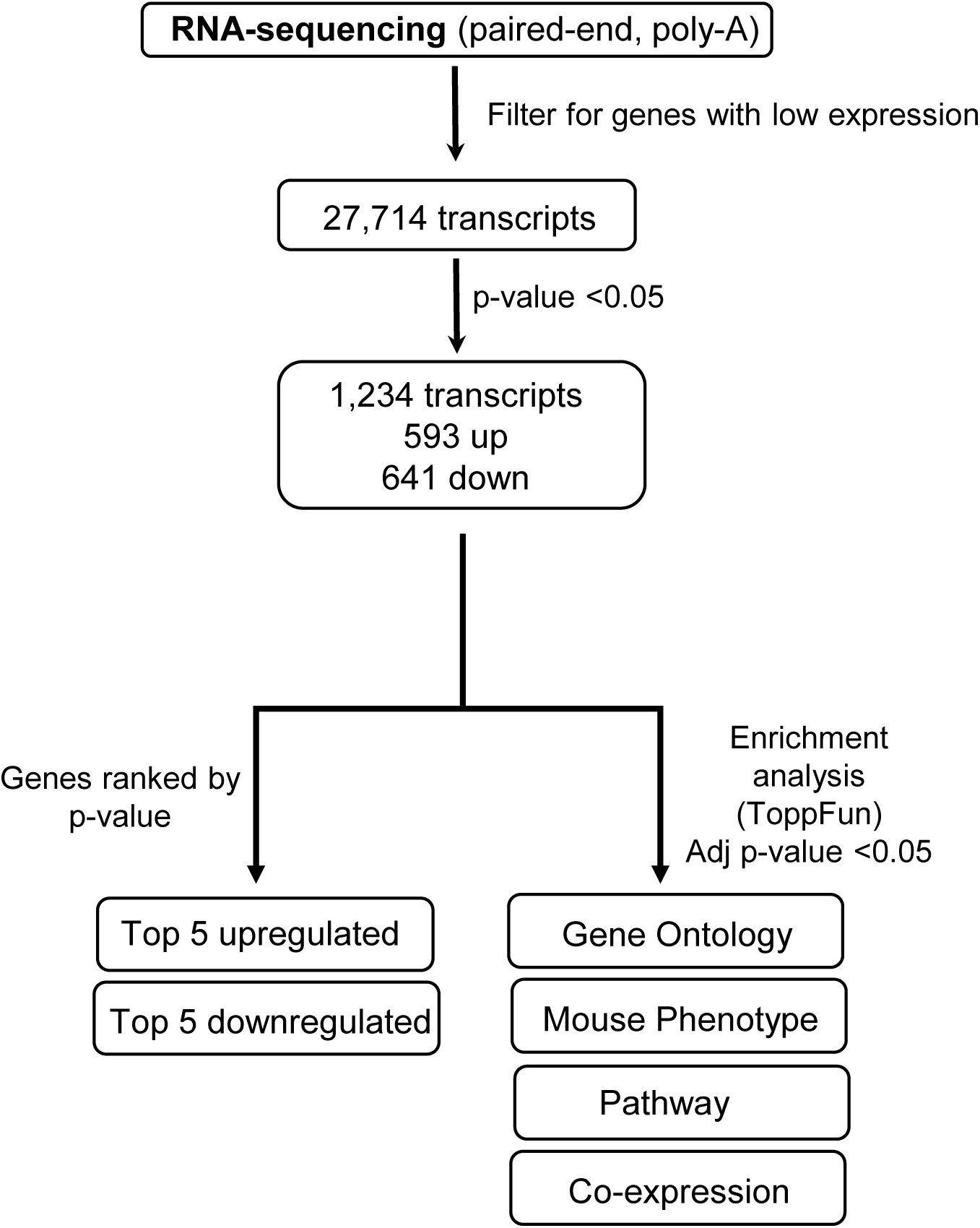
RNA sequencing methodological workflow after TG treatment. Preliminary filtering for genes with low expression and p-value cutoff of 0.05 was used to uncover differentially expressed genes. ToppFunn application of the ToppGene Suite was used to to reveal pathways of interest using padj <0.05 = significant.

**Figure 4:**
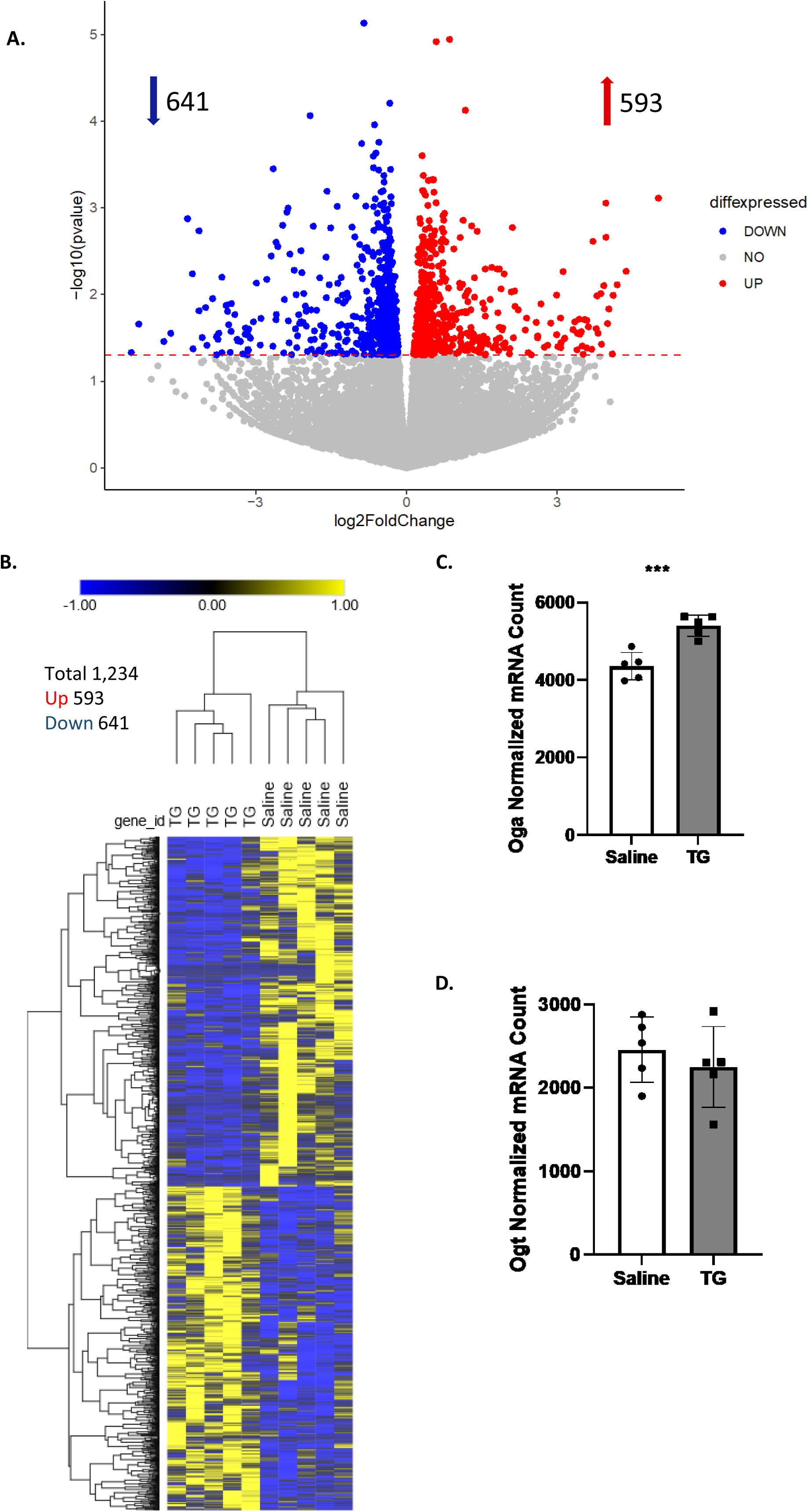
Differential gene expression in response to TG treatment. **A)** Volcano plot of all differentially expressed genes after TG treatment. A total of 27,714 expressed genes were detected. Using a p-value threshold of <0.05 for the difference between TG and saline, 593 genes were upregulated and 641 genes were downregulated. Upregulated genes are represented in red and downregulated genes in blue. Genes that did not reach the p-value threshold of <0.05 are shown in gray. **B)** Significant differentially expressed genes were also represented as a heatmap with hierarchal clustering (n=1,234). Each column on the x-axis represents a sample and each row on the y-axis represents a gene’s z-score. The dendrogram above the heatmap shows the hierarchal clustering. Genes with high expression are shown in yellow while genes with low expression are in blue. **C)** OGA mRNA normalized count in the TG compared to saline treated mice. **D)** OGT mRNA normalized count was unchanged in TG compared to saline treated mice. ***p < 0.001. n=5 mice per group.

Using these data, our next level of analysis was directed at determining the top 5 up and downregulated genes. The list of genes, fold changes, and p-values are listed in **Table 1**. As mentioned above, OGA mRNA is increased and features as the top upregulated gene (**Figure 4C**). Cell division/development are potential targets for TG treatment since the top 2 upregulated genes are Cep95 (centrosomal protein 95) and Ccar1 (cell division cycle and apoptosis regulator 1) and Sox18 (SRY-Box transcription factor 18) which is involved in embryonic development was down regulated. Changes in genes related to metabolism were a consistent feature of the top regulated mRNAs. Specifically, Pdhx (pyruvate dehydrogenase complex component X) was upregulated and Fads2 (fatty acid desaturase 2) was down regulated consistent with metabolic switching involving unsaturated fatty acids and the TCA cycle. Interestingly, Il20rb (interleukin 20 receptor subunit beta) was a strong upregulated gene consistent with modulating of inflammatory signaling. Three cell structural genes were among the top downregulated genes including nerve cell intermediate filament protein Nes (Nestin), an essential component of the plasma membrane Tmem184a (transmembrane protein 184A) and collagen chaperone Serpinh1 (serpin family H member 1).

**Table 1:**
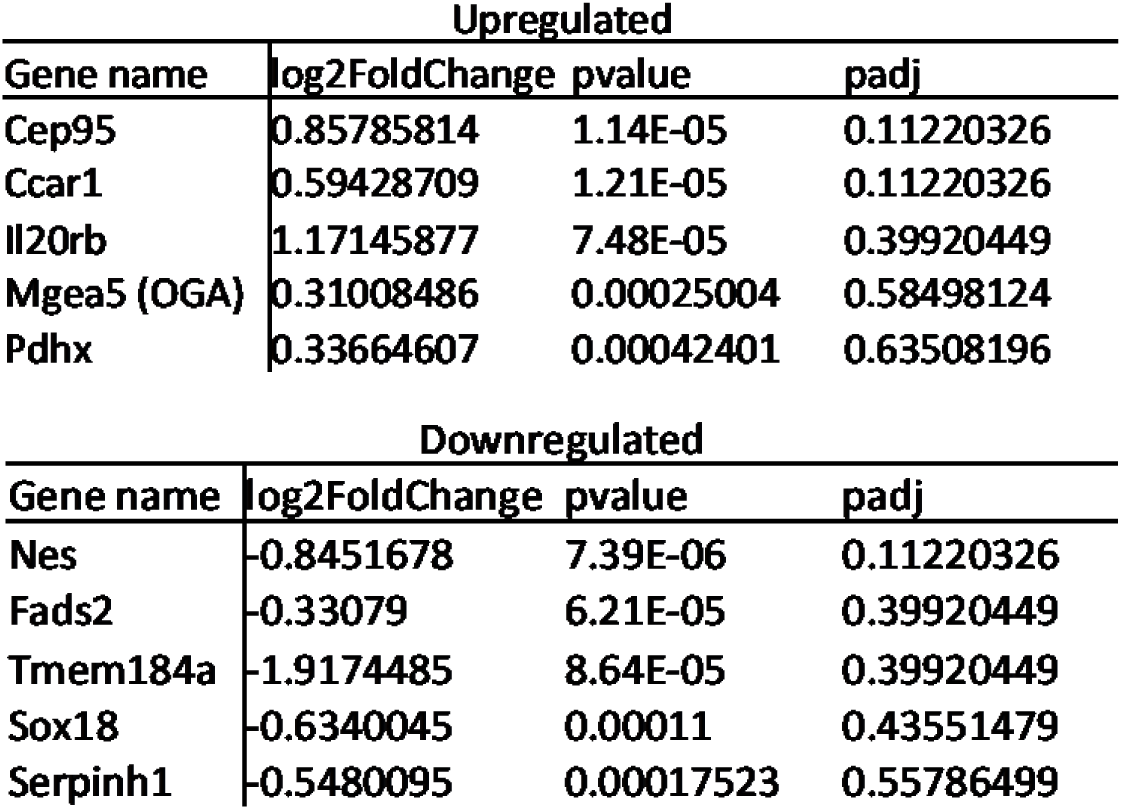
Top 5 upregulated and downregulated genes in response to 3 hours of TG treatment ranked by p value.

### Functional enrichment pathway analysis

To identify the key biological domains that respond to TG treatment, we performed functional enrichment analysis using all 1,234 detected transcripts. The analysis revealed 481 upregulated and 1,617 downregulated pathways in response to TG (**Supplemental Table 5**). From these, we selected notable pathways of interest previously implicated in neurodegenerative diseases for a more detailed representation in network maps. A network representation of the upregulated pathways of interest is shown in **Figure 5A**, which shows 5 rectangular nodes representing different pathways and genes in oval nodes. One upregulated pathway of interest was “Alzheimer’s Down”, which contained genes that are typically down in Alzheimer’s disease. **Figure 5B** shows bar graphs for genes in the Alzheimer’s down pathway, including Pdhx (Log2Fold Change= 0.3366 p=0.0004), Sucla2 (Succinate-CoA Ligase ADP-Forming Subunit Beta; Log2Fold Change= 0.2517 p=0.0030), and Ndufs4 (NADH:Ubiquinone Oxidoreductase Subunit S4; Log2Fold Change= 0.3181 p=0.0486). Similar to the gene level analysis, many genes in this pathway were related to metabolism. Another upregulated pathway of interest was the centrosome pathway. The centrosome pathway was composed of transcripts such as Cep95 (Log2Fold Change= 0.8579 p=1.14E-05), Ccnb2 (Cyclin B2; Log2Fold Change= 1.7816 p=0.0379), and Plekha7 (Pleckstrin Homology Domain Containing A7; Log2Fold Change= 0.3401 p=0.0024), as shown in **Figure 5C**. Again, in line with the gene level analysis, this pathway contained genes related to cell division. The full gene list for all pathways of interest is presented in **Supplemental Table 6**.

**Figure 5:**
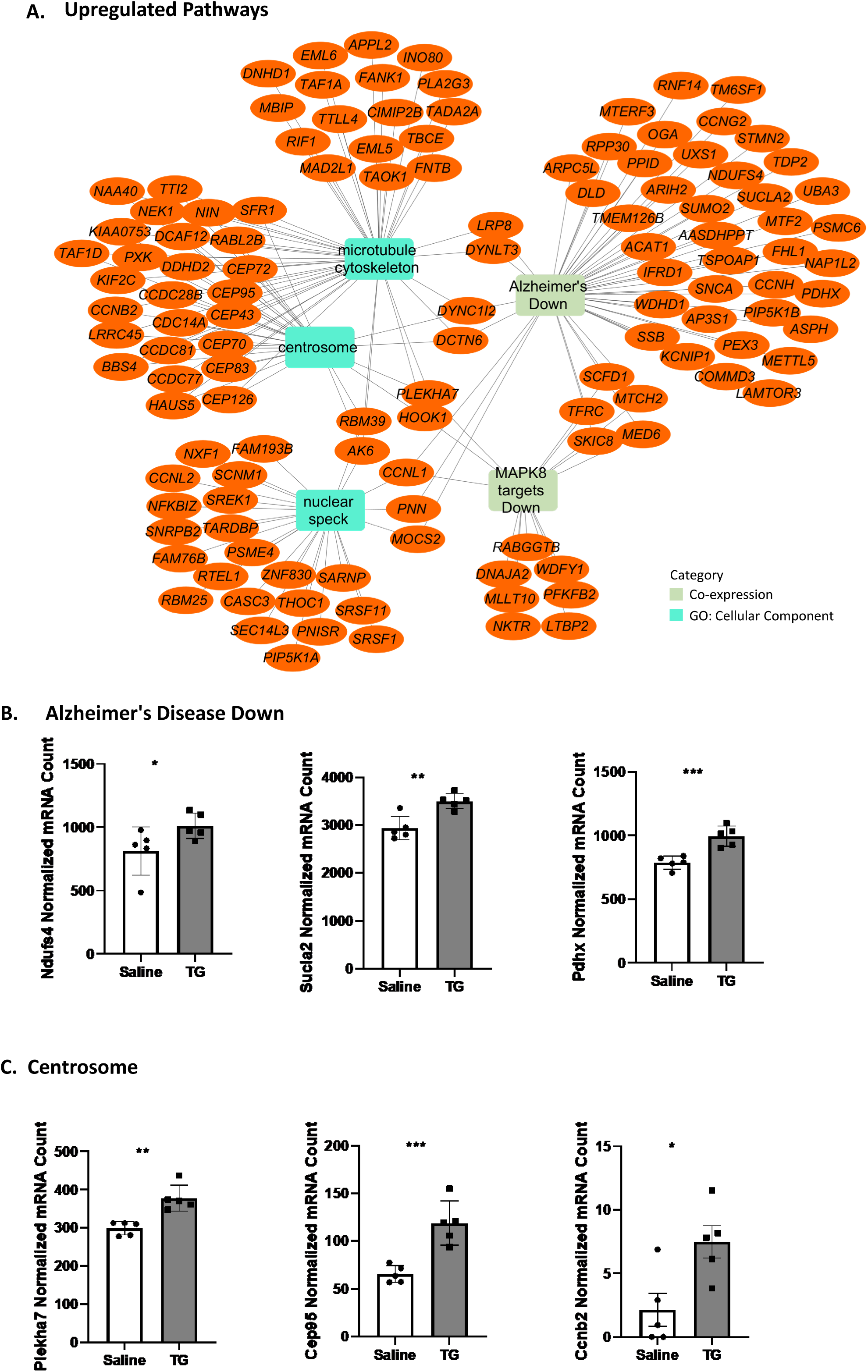
Representation of selected upregulated pathways from enrichment analysis. **A)** Representative network map of selected pathways of interest with accompanying genes. Orange nodes are genes upregulated following TG treatment while the rectangular nodes are enriched pathways. Functional enrichment was performed using theToppFun application of the ToppGene Suite. Network was generated using Cytoscape. Different colors of the rectangle represent the different pathway category shown in the legend. **B)** Bar graphs showing the normalized mRNA count of representative genes in the Alzheimer’s Disease Down pathway. Genes include Ndufs4 (NADH:Ubiquinone Oxidoreductase Subunit S4), Scula2 (Succinate-CoA Ligase ADP-Forming Subunit Beta), and Pdhx (Pyruvate dehydrogenase complex component X). **C)** Bar graphs of the normalized mRNA count for top genes of the centrosome pathway: Plekha7 (Pleckstrin Homology Domain Containing A7), Cep95 (centrosomal protein 95), and Ccnb2 (Cyclin B2). *p < 0.05, **p < 0.01, ***p < 0.001. n=5 mice per group.

Downregulated pathway analysis can be found in **Figure 6**. **Figure 6A** shows a network representation of downregulated pathways of interest, with pathways being in 16 rectangular nodes and genes in oval nodes. Downregulated pathways of interest include many cell structural pathways, such as cell adhesion, actin filament binding, and extracellular matrix. This is consistent with the top down regulated genes discussed above being related to cell structure. Additionally, genes that are normally upregulated in Alzheimer’s disease and aging were found to be downregulated, inverse to what was observed in the upregulated analysis. **Figure 6B** shows bar graphs for genes upregulated in Alzheimer’ s disease such as Gfap (Glial Fibrillary Acidic Protein; Log2Fold Change= -0.2328 p=0.0376), Vcam1 (Vascular Cell Adhesion Molecule 1; Log2Fold Change= -0.2515 p=0.0153), and Adgrg1 (Adhesion G Protein-Coupled Receptor G1; Log2Fold Change= -0.2493 p=0.0146). Representative bar graphs for genes normally up in aging are in **Figure 6C**, specifically Gsn (Gelsolin; Log2Fold Change= -0.3083 p=0.0007), Fgfr2 (Fibroblast Growth Factor Receptor 2; Log2Fold Change= -0.2989 p=0.0009), and Abcg1 (ATP Binding Cassette Subfamily G Member 1; Log2Fold Change= -0.3128 p=0.04997). To represent the multiple cell structural pathways measured, **Figure 6D** shows genes of the cell adhesion pathway, such as Actn1 (Actinin Alpha 1; Log2Fold Change= -0.1946 p=0.0374), Lamb2 (Laminin Subunit Beta 2; Log2Fold Change= -0.4447 p=0.0011), and Itgb5 (Integrin Subunit Beta 5; Log2Fold Change= -0.3045 p=0.0081). All genes for each pathway can be found in **Supplemental Table 6.**

**Figure 6:**
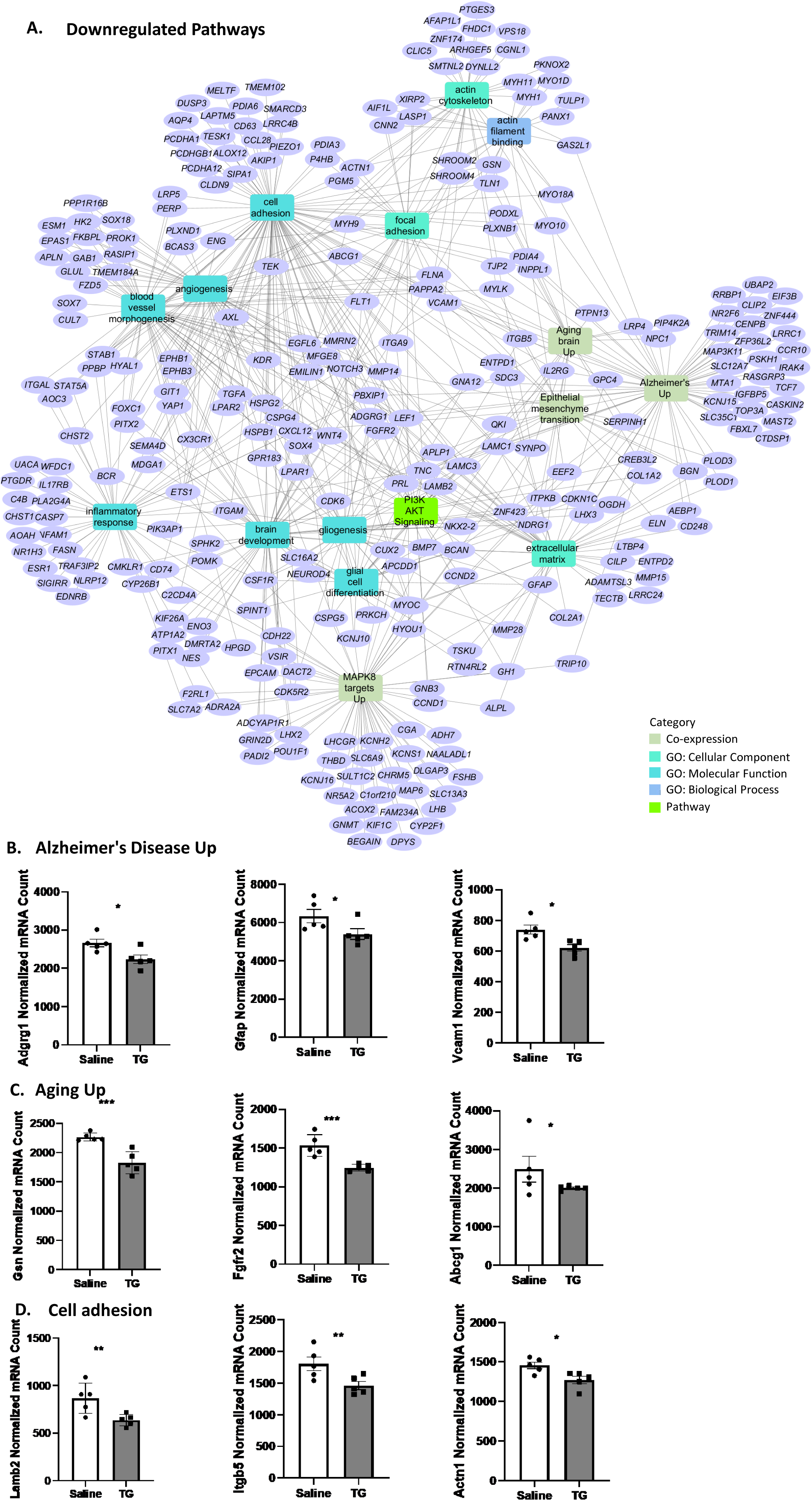
Representation of selected downregulated pathways from enrichment analysis. **A)** Network map of selected downregulated pathways of interest with associated genes. Purple nodes are genes downregulated following treatment while the rectangular nodes are enriched pathways. Functional enrichment was performed using ToppFun application of the ToppGene Suite and the network was generated using Cytoscape. Different rectangle colors represent the different pathway category shown in the legend. **B)** Bar graphs showing the normalized mRNA count of representative genes of the Alzheimer’s Disease Up pathway. Genes include Adgrg1 (Adhesion G Protein-Coupled Receptor G1), Gfap (Glial Fibrillary Acidic Protein), Vcam1 (Vascular Cell Adhesion Molecule 1). **C)** Bar graphs of the normalized mRNA count for top genes of the Aging Up pathway: Gsn (Gelsolin), Fgfr2 (Fibroblast Growth Factor Receptor 2) and Abcg1 (ATP Binding Cassette Subfamily G Member 1). **D)** Bar graphs of the top genes of the cell adhesion pathway normalized mRNA count: Lamb2 (Laminin Subunit Beta 2), Itgb5 (Integrin Subunit Beta 5), and Actn1 (Actinin Alpha 1). *p < 0.05, **p < 0.01, ***p < 0.001. n=5 mice per group.

### Comparison of differentially expressed acute and chronic genes

To assess early versus late gene expression in response to elevated O-GlcNAcylation, we compared differentially expressed genes between acute (this study) and chronic TG treatment (as we previously published^30^) (**Figure 7**). For acute treatment, we used 1,234 significantly differentially expressed genes from the dataset introduced in this manuscript (**Supplemental Table 4**). 7 genes in the acute dataset were novel and unnamed; therefore, they were not used for this analysis giving the total number of genes for this comparison 1,227. For chronic treatment, we used transcriptomic data from a previous manuscript^30^. In brief, chronic TG treatment (35 mg/kg) was administered in 1-month old mice biweekly for 2.5 months (n=12 each group). The brain was harvested, and transcriptomic analysis revealed 4,145 significant genes after chronic TG treatment. The gene list for acute and chronic treatment can be found in **Supplemental Table 7.**

**Figure 7:**
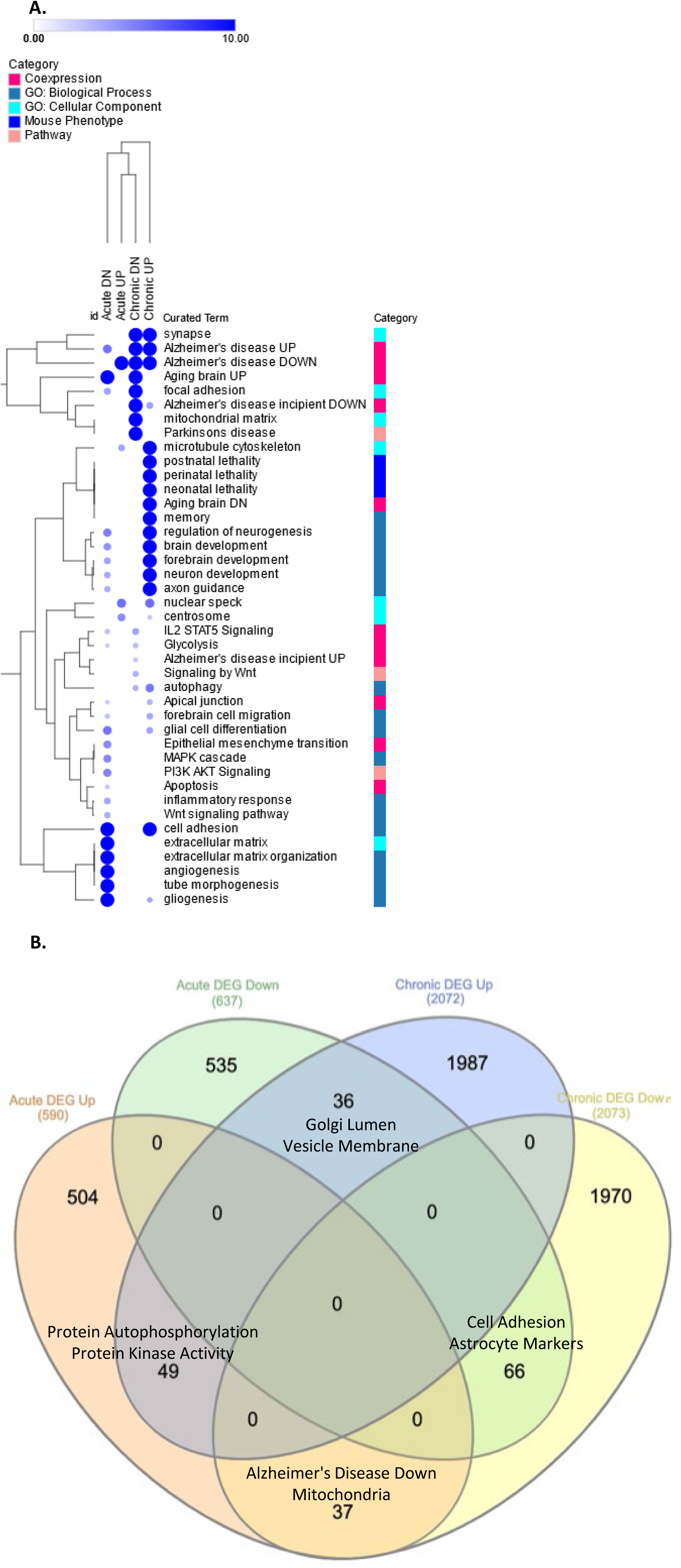
Comparison of enriched pathways and differentially expressed genes (DEG) found after acute and chronic Thiamet G (TG) treatment. Significant differential expressed genes of the acute dataset introduced in this manuscript were compared to RNAseq data from a previously published manuscript investigating chronic (2.5 month) treatment of TG^30^. **A)** Heatmap representation of select pathways enriched in genes differentially expressed in acute and chronic treatment of TG. Functional enrichment was performed using ToppCluster application. Heatmap was generated using Morpheus application. The size and color of the circles in the heatmap are proportional to the significance of enrichment (negative log of p-value FDR). Hierarchical clustering of rows and columns was done using Euclidean distance and complete linkage. **B)** Venn diagram of the intersecting genes after acute and chronic TG. ToppFun was used to identify the pathways of intersecting genes. n=5 mice per group for acute treatment. n=12 mice per group for chronic treatment.

The first step of the analysis was to compare the upregulated and downregulated pathways represented from the acute and chronic genes (**Figure 7A**). Pathways that were upregulated in both acute and chronic datasets were microtubule cytoskeleton and nuclear speck. In contrast, pathways that were down in both datasets were focal adhesion and genes up in aging. Gene list for all pathways in the heatmap can be found in **Supplemental Table 8.**

Next, we compared intersecting genes of the two datasets and found 49 genes were up in both acute and chronic TG treatment and 66 genes were down in acute and chronic TG treatment (**Figure 7B and Supplemental Table 9**). Pathway analysis revealed that genes up in both acute and chronic TG treatment were related to protein autophosphorylation and kinase activity (**Figure 7B**), including Eif2ak4 (Eukaryotic Translation Initiation Factor 2 Alpha Kinase 4), Prkab2 (Protein Kinase AMP-Activated Non-Catalytic Subunit Beta 2), and Trpm7 (Transient Receptor Potential Cation Channel Subfamily M Member 7) (**Supplemental Table 10**). 66 downregulated genes in acute and chronic datasets were related to cell adhesion and astrocyte markers (**Figure 7B and Supplemental Table 9**).These pathways included genes such as Vcam1 and Gfap (**Supplemental Table 10**).

On the other hand, there were 37 genes that were up after acute treatment and down after chronic treatment (**Figure 7B and Supplemental Table 9**). Pathway analysis showed these genes were down in Alzheimer’s disease and related to mitochondria (**Figure 7B**), and including mitochondria-related transcripts such as Ndufs1 (NADH Dehydrogenase subunit 1), Ndufs4 (NADH Dehydrogenase subunit 1), and Mtch2 (Mitochondrial Carrier 2) (**Supplemental Table 10**). There were also 36 genes that were down after acute treatment and up after chronic treatment, which were related to the Golgi lumen and vesicle membrane (**Figure 7B and Supplemental Table 9**). This included genes such as Tgfa (Transforming Growth Factor Alpha) and Wnt4 (Wnt Family Member 4) (**Supplemental Table 10**).

## DISCUSSION

In this manuscript, we performed transcriptome analysis to examine the acute effects of TG treatment in mouse brain to identify transcripts or pathways that are influenced by elevation of protein O-GlcNAcylation. We found that after 3 hours of TG treatment overall O-GlcNAc protein expression was increased as expected, whereas no changes were found in OGA and OGT protein levels. Therefore, using this treatment paradigm we were able to determine acute effect of TG treatment, before any obvious compensation in OGA and OGT protein levels as previously observed in chronic TG treatment^30,37^. RNA sequencing analysis revealed 1,234 signficant genes that were significant after 3 hours of TG treatment. This supports our hypothesis that acute TG treatment induces significant changes to the mouse transcriptome in as early as 3 hours. When focusing on the mRNA expression of the enzymes that control O-GlcNAc abundance, OGA and OGT, we observed a significant increase in OGA mRNA and no change in OGT. This suggests that OGA mRNA increased to compensate or counteract the pharmacological inhibition of OGA by TG in as soon as 3 hours. The observed increase in OGA mRNA and unchanged OGA protein expression is consistent with previous data from chronic TG treatment^30^.

### Changes in cell division and structure

Acute TG treatment revealed that the top two upregulated genes were related to cell division and upregulated pathway of interest was centrosome. This is consistent with the literature showing that O-GlcNAc emerging as a regulator of cell cycle progression^38^. In particular, the top upregulated gene Cep95 is found in the centrosome and spindle pole^39^. The centrosome and spindle poles are important parts of mitosis or the M phase of the cell cycle. This stage of the cell cycle involves the formation of the mitotic spindles and separating duplicated chromosomes^40^. It has been shown that dysregulation (increase or decrease) of O-GlcNAc by modifying OGA or OGT causes disruption in mitosis that leads to defects in assembly of mitotic spindles as well as delayed M phase progression^41–43^. In addition, Ccar1 was the second highest upregulated gene. Ccar1 is also involved in apoptosis signaling and cell cycle regulation by enabling RNA polymerase II to bind to DNA^39,44^. This is interesting due to the known role of O-GlcNAc to modify RNA polymerase II and other transcription factors to regulate gene expression^30,31,45^. Furthermore, 29 peptides of Ccar1 are known to contain O-GlcNAc sites^46^. Additionally, studies show there is cell cycle dysregulation in AD^47^ suggesting OGA inhibitors could impact AD pathogenesis or progression.

Acute TG treatment revealed downregulated genes related to cell structure accompanied by downregulated pathways related to cell adhesion. One top downregulated gene is Nes, which is an intermediate filament protein expressed in nerve cells^48^. Previous literature has shown the O-GlcNAc modification of intermediate filament proteins such as keratin^49^ and neurofilament (H, M and L)^50^, however no work has been done thus far on the relationship between O-GlcNAc and Nes. Other genes related to cell adhesion and structure were measured, including multiple isoforms of actin, laminin and integrin. It has been shown that cell adhesion molecules are dysregulated in AD; however, the exact mechanism is unclear^51–53^. Downregulation of the cell adhesion pathway was also measured in overlapping genes in response to acute and chronic TG treatment, giving more reason to further investigate the role of O-GlcNAc in aspects of cell structure and remodeling.

### Impact on cell signaling

One downregulated pathway of interest was PI3K-Akt signaling. A previous study from our lab showed mTOR and mTOR related transcripts were upregulated in response to chronic (2.5 months) TG treatment. This is interesting due to mTOR being downstream of PI3K-Akt. PI3K-Akt-mTOR signaling has been shown to be suppressed as Aβ is increased in a mouse model of AD^54^. In addition, p-mTOR was increased in AD postmortem brain and colocalized with hyperphosphorylated tau^55^. Interestingly, there is also a known association of O-GlcNAc abundance and PI3K-Akt signaling in various neurodegenerative disease^56,57^. Previous literature also showed a negative correlation between O-GlcNAc and Akt phosphorylation after 24 hours of TG treatment in male mice^58^. Because this study is after only 3 hours, we hypothesize PI3K-Akt signaling is regulated by O-GlcNAc abundance, and more experiments should be done to further understand this process.

Additional signaling pathways emerged when comparing the transcriptome of acute and chronic TG mice brain to reveal pathways influenced by O-GlcNAc. This comparison revealed transcripts related to protein autophosphorylation and protein kinase activity were upregulated in response to acute and chronic TG treatment. The gene that encodes for the regulatory subunit of the AMP-activated protein kinase (AMPK)^59^, Prkab2, was seen in the protein kinase activity pathway. AMPK hyperphosphorylation has been linked to pathological features of AD^60^ and can directly phosphorylate tau, leading to neurofibrillary tangles^61^. Additionally, Trpm7 functions as a Ca^2+^and Mg^2+^channel and has been shown to exhibit decreased activities in various neurodegenerative diseases^62^. In AD, Trpm7 is suggested to contribute to Aβ pathology^63^. How O-GlcNAc regulates Trpm7 in neurodegenerative diseases will be an interesting future direction.

### Dysregulated metabolism

One of the most interesting findings of this study was the changes in metabolism and AD-related pathways. Dysregulation of metabolism is a key pathological feature of AD^64–66^, and it can be directly manipulated by increasing O-GlcNAc abundance. One of the top upregulated genes following acute TG treatment was Pdhx, a component of pyruvate dehydrogenase responsible for linking glycolysis to the TCA cycle^67,68^. Pyruvate dehydrogenase is decreased in the parietal and temporal cortex of AD patients^69^. Pathway analysis revealed that genes that are normally down in AD, such as metabolism and signaling genes^70^, are up with acute TG treatment. Furthermore, genes normally upregulated in AD and aging, such as structural genes and astrocyte markers^70,71^, are down with acute TG treatment. These changes measured at baseline in wildtype mice supports the use of OGA inhibitors as AD treatment.

Conversely, a different pattern emerged when comparing the intersecting genes of acute and chronic TG-treated mice brains. In agreement with the previous paragraph, genes that are normally down in AD are up after acute TG; however, in chronic TG these genes are down similar to what we observe in AD. Intersecting genes included Ndufs1 and Ndufs4, which are both subunits of complex 1 of the electron transport chain^72^. Complex I dysfunction has been reported in AD^73^ and other neurodegenerative disorders^74^. Additionally, mutations in Ndufs1 lead to increased H2O ^75^, and mutations in Ndufs4 prevent complex I assembly^75,76^. Mutations in either subunit led to a decrease in complex activity^76,77^. This leads to a conflicting theory regarding the long-term effects of OGA inhibitors for AD treatment because the downstream effects are possibly harmful. More longitudal studies should be done to investigate the use of OGA inhibitors in AD.

### Limitations and Conclusions

There are limitations in this study that must be noted. In this experiment, only male mice were used; therefore, females mice will be needed in future studies. We also used one time-of-time for our experiment, and future studies with different time-of-day can be informative as circadian regulation of gene expression can impact the effects of protein O-GlcNAcylation^78–80^.

From this study, we determined the mouse brain transcriptome after 3 hours of elevating protein O-GlcNAcylation using the OGA inhibitor TG. These data provide insights into the mechanism of action of this dynamic post-translational modification. We identified 1,234 significant transcripts that were differentially expressed. Pathway enrichment analysis revealed upregulated genes related to a decrease in AD and downregulated genes that are normally increase in AD. However, comparison of acute and chronic TG treatment revealed genes that are normally down in AD are down after chronic TG treatment. This led us to conclude that OGA inhibitors are promising for the treatment of AD; however, their downstream chronic effects may be detrimental with regard to mitochondrial function.

### Ethics approval and consent to participate

Animal study performed in this manuscript has been approved by UAB IACUC. No human studies were conducted.

## Availability of data and materials

Please contact the author to request data.

## Author contributions

MB and MSK performed experiments and participated in data analyses. MB, XO, MSK, JCC, VDU, and JZ participated in data analyses and interpretation. MB, VDU, and JZ wrote the manuscript. All authors edited and approved the final manuscript.

## Acknowledgments

We thank members of Drs. Chatham, Darley-Usmar, and Zhang laboratories for discussions.

